# Designing pathways for bioproducing complex chemicals by combining tools for pathway extraction and ranking

**DOI:** 10.1101/2024.06.10.598209

**Authors:** Anastasia Sveshnikova, Omid Oftadeh, Vassily Hatzimanikatis

## Abstract

The synthesis of many important biochemicals involves complex molecules and many reactions. Therefore, the design and optimization of whole-cell biocatalysts to produce these molecules requires the use of metabolic modeling. Such modeling involves the extraction of the production pathways from biochemical databases and their integration into genome-scale metabolic models of the host organism. However, the synthesis of such complex molecules requires reactions from multiple pathways operating in balanced subnetworks that are not assembled in existing databases. Here we present SubNetX, a novel computational algorithm that extracts reactions from a given reaction database and assembles balanced reaction subnetworks to produce a target biochemical from a selected set of precursor metabolites, energy currencies, and cofactors of the host organism. These subnetworks can be directly integrated into whole-cell metabolic models, and using available methods, we can then reconstruct all alternative biosynthetic pathways and rank them according to design criteria such as yield, pathway length, and other optimization goals. We applied SubNetX to eight selected secondary metabolites and one non-natural chemical used as pharmaceuticals to demonstrate the potential of this pipeline.

## Introduction

The green chemistry movement is shifting the production of important chemicals, such as pharmaceuticals or food additives^1^, away from traditional fossil–fuel-based chemical syntheses and costly agricultural extraction towards bioproduction^2^. Eco-friendly and sustainable bioproduction uses microbes to produce chemicals of interest, either taking advantage of existing biosynthetic pathways or through metabolic engineering^3^. However, the complexity of biochemicals often limits their industrial scalability, and engineering strategies are currently limited to relatively simple compounds (e.g., ethanol and 1,3-butanol)^4,5^. One reason for this is that engineering strategies mainly propose linear pathways in contrast with living organisms that can combine metabolic pathways to generate complex secondary metabolites. Inspired by nature, researchers thus aim to design pathways that divert resources from several pathways toward a single target to achieve higher yields for complex natural and non-natural compounds.

Engineering strategies rely on biochemical databases to search and suggest pathways. Some biochemical databases contain the natural reactions observed in nature^6,7^, while others contain computationally predicted reactions that attempt to represent the conceivable biochemical space^8,9^. Different computational algorithms are used to search these databases to find bioproduction pathways to a particular target compound. Currently, bioproduction pathways are designed using three classes of computational tools (for a comprehensive review, see Wang et al.^10^): (i) graph-based approaches that use graph search algorithms to find pathways, (ii) stoichiometric approaches that use constraint-based optimization to find pathways, and (iii) retrobiosynthesis approaches that use algebraic operations to propose novel reactions. Graph-based and retrobiosynthesis methods both rely on graph-search algorithms, enabling them to navigate large networks of biochemical reactions. However, these pathways are a linear combination of heterologous reactions limiting the output of such methods to pathways with a single precursor among the host metabolites^10^. However, a pathway is not stoichiometrically feasible if the required cosubstrates and cofactors are not connected to the host metabolism, a potential shortcoming of linear pathways. On the other hand, stoichiometric approaches allow for the analysis of subnetworks connected to the host metabolism via multiple precursors and their evaluation in the context of the host metabolism. As such, they often yield feasible pathways. However, constraint-based approaches are sensitive to the size of the reaction network due to limited computational power and cannot account for conceivable xenobiotic metabolism that contains hundreds of thousands of reactions^8,11^ ^12,13^.

Here, we address issues in existing pathway-design tools by combining the strengths of constraint-based and retrobiosynthesis methods into a novel pipeline, termed SubNetX (for **Subnet**work e**x**traction). This pipeline allows the exploration of large reaction networks to find an optimal pathway for the bioproduction of a target compound that would integrate into the native host metabolism, while accounting for the stoichiometric and thermodynamic feasibility of the pathways. SubNetX relies on constraint-based methods to ensure the feasibility and retrobiosynthesis methods to handle larger networks and identify novel biosynthetic pathways. We show that our approach predicts viable pathways for the synthesis of eight chemical targets with higher production yields compared to linear pathways. We also demonstrated the potential of our method in a case study designing novel biosynthetic pathways in *E. coli* for tadalafil, a therapeutically relevant compound. The pipeline presented here is readily transferable to other biochemicals, and we discuss how it can be applied to other host organisms. Thus, we anticipate that SubNetX will be instrumental in designing pathways producing complex natural and non-natural compounds in various organisms.

## Results and Discussion

### SubNetX transforms large networks of predicted reactions into thermodynamically feasible biochemical pathway designs

To maximize bioproduction strategies, we designed SubNetX to employ network exploration tools and constraint-based optimization methods to link required cosubstrates and subsequent byproducts to the host’s native metabolism (Supplementary Fig. 1). The SubNetX workflow and its application to the prediction of balanced minimal subnetworks is illustrated in Figure 1. The SubNetX workflow is organized into five main steps: (i) reaction network preparation where a database of elementally balanced reactions, target compounds, and precursor compounds are defined; (ii) graph search of linear core pathways from the precursor compounds to the target compounds, (iii) expansion and extraction of a balanced subnetwork where cosubstrates and byproducts are linked to the native metabolism, (iv) integration of the subnetwork into the host, and (v) ranking of the feasible pathways. Overall, the workflow requires as inputs: (i) a network of balanced biochemical reactions, i.e., a database, (ii) a set of target compounds, (iii) a set of precursors, which most commonly depend on the host of choice, (iv) a metabolic model of the host metabolism, and (v) user-defined parameters to adjust the search based on the user’s needs. The network of balanced biochemical reactions can be either limited to known reactions or extended to include predicted biochemical reactions.

**Figure 1:**
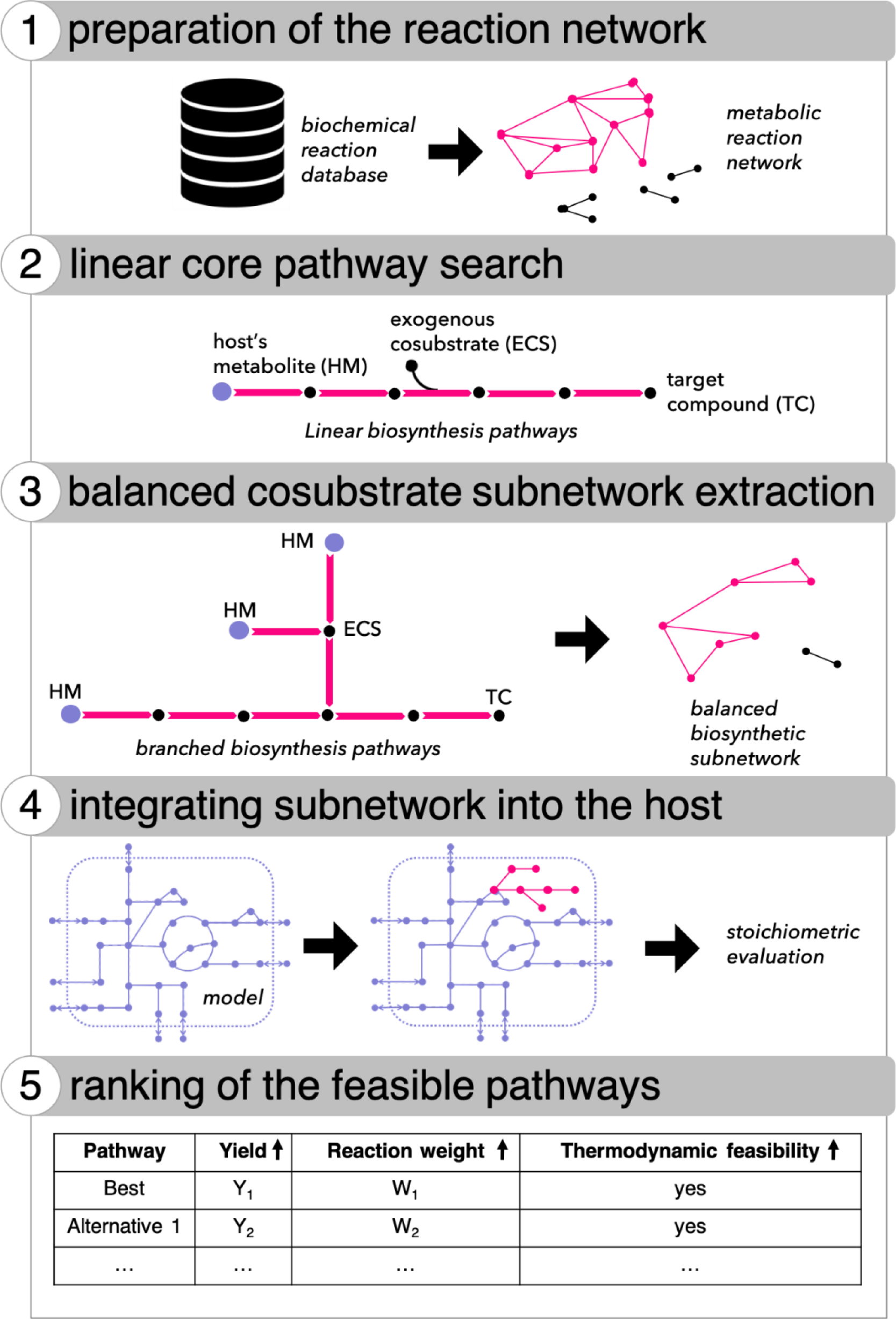
The SubNetX workflow. Step 1: preparation of the reaction network; step 2: linear core pathway search; step 3: balanced cosubstrate subnetwork extraction; step 4: integrating subnetwork into the host; and step 5: ranking feasible pathways.

To find pathways to the targets and required cosubstrates in this work, we prepared a network of known and predicted biochemical reactions designed for the biosynthesis of aromatic compounds^13^ (Fig. 1, steps 1, 2, and 3). We then integrated the resulting subnetwork into the genome-scale metabolic model of *E. coli* (Fig. 1, step 4) to ensure that the target compound can be produced according to the metabolic capabilities of the host. As the extracted networks could contain thousands of reactions, it would be experimentally impossible to integrate the entire network into the host. Therefore, we used a mixed-integer linear programming (MILP) algorithm to identify sets of feasible pathways that will contain a much smaller number of heterologous steps (Fig. 1, step 5). We did this by finding the minimum number of essential reactions from the subnetwork that could produce the target compound, with each minimal set of reactions referred to as a feasible pathway. Finally, these feasible pathways were ranked based on yield, enzyme specificity, and thermodynamic feasibility (Fig. 1, step 5).

### SubNetX successfully predicts balanced biosynthesis subnetworks for complex secondary metabolites

As an initial test of SubNetX, we applied it to eight selected secondary metabolites used as pharmaceuticals: alkaloids (berberine, ajmalicine, scopolamine, strictosidine), products of the shikimate pathway (N-cinnamoyl serotonin, benzyl benzoate, benzyl cinnamate), and glycosylated isoflavonoid (quercetin 3-O-[6’-acetyl-glucoside]). We evaluated the synthetic accessibility of these compounds^14^ (Methods: Selection of the case study compounds; Supplementary Table 1) among a list of various natural compounds synthesized in different hosts (Supplementary Table 2). We searched the ARBRE network^13^ to connect the target molecules to the host metabolism. For each compound, we extracted a balanced subnetwork as described in Methods (Figure 2).

**Figure 2:**
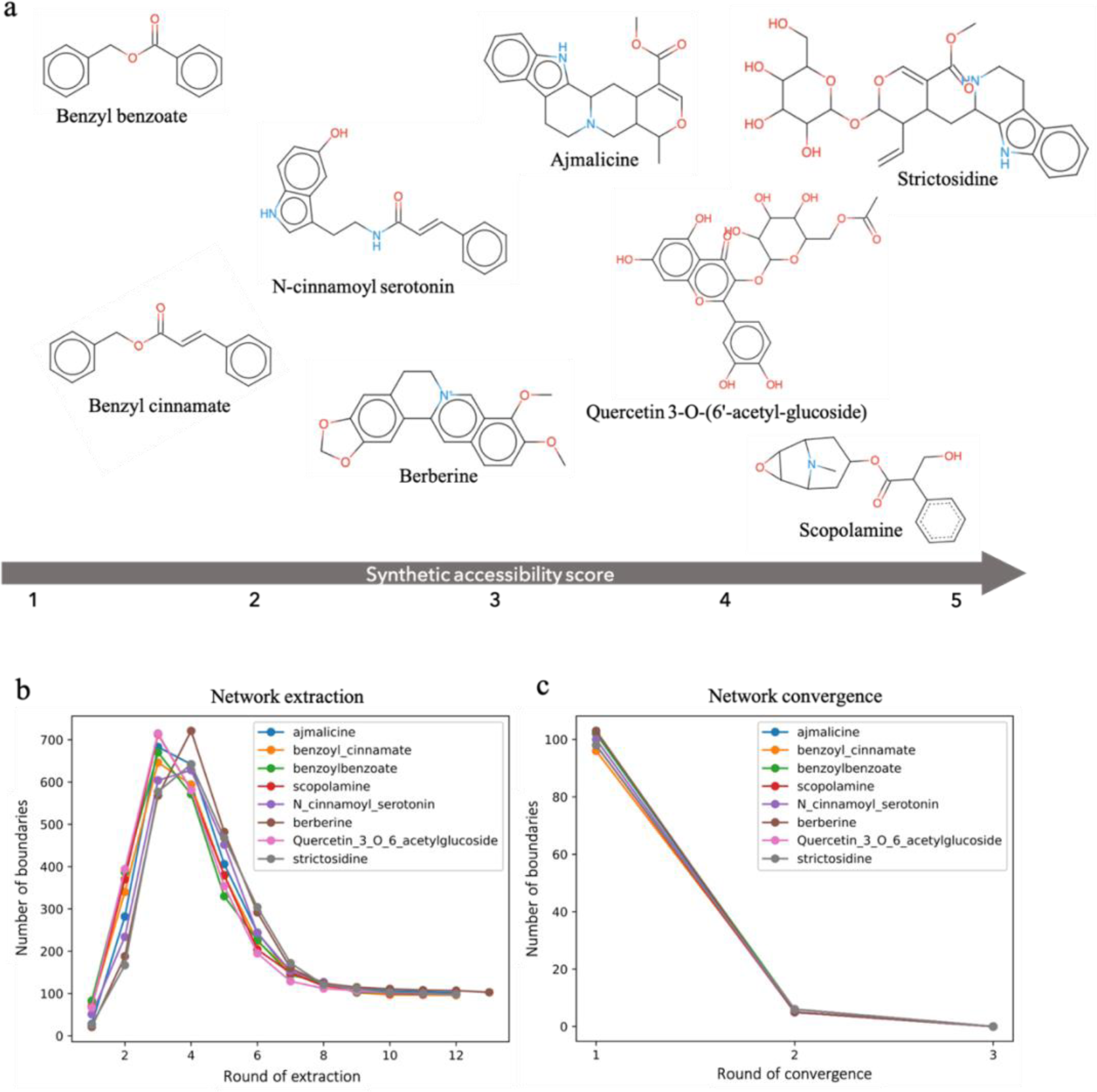
Subnetwork extraction for the case studies. **a** Structure of each case study compound corresponding to the synthetic accessibility score. **b** Network extraction statistics showing the number of boundaries in each round of subnetwork extraction. **c** Network convergence statistics showing the number of boundaries in each round of subnetwork convergence.

Using only the ARBRE biochemical network, we mapped seven of the target compounds to *E. coli* native metabolites. For the eighth, scopolamine, the ARBRE biochemical network did not contain the biosynthesis pathway, producing two tropane derivatives from putrescine, which is essential for scopolamine production (Supplementary Fig. 2). Thus, we supplemented these pathways using ATLASx^11^, which can fill in missing reactions to connect scopolamine to the *E. coli* metabolism. ATLASx recovered a pathway to produce the two tropane derivatives already used to experimentally produce scopolamine in *E. coli*^15^. This pathway included one unbalanced reaction converting N-methylpyrrolinium to tropinone, which we replaced by two balanced reactions, chalcone synthase and tropinone synthase (Supplementary Fig. 2). These two reactions were annotated and added to ARBRE, which allowed us to create a balanced subnetwork for scopolamine. The synthesis of scopolamine provides an example of how potential gaps in biochemical knowledge can be identified and addressed while designing pathways to produce novel compounds.

Once we were able to extract subnetworks for all eight compounds, we compared the sizes of the extracted subnetworks, finding them to be similar for different targets (Figure 2). Interestingly, alternative feasible pathways required network expansion for the balanced synthesis of cofactors such as tetrahydrobiopterin that are only found in vertebrates^16^. Such non-native cofactors were only necessary for some pathways of the extracted subnetworks, while alternative feasible pathways existed involving only *E. coli* cofactors. Thus, to control network size and avoid unnecessary expansion, we recommend using a SubNetX search mode that avoids expansion around non-native cofactors.

### Constraint-based optimization extracts feasible branched pathways

Next, we evaluated the stoichiometric feasibility of the extracted subnetworks, which were designed to biosynthesize each compound using constraint-based optimization. To this end, we integrated the subnetwork for each compound into the *E. coli* genome-scale metabolic model iJO1366^17^ and maximized the production of the target compounds. We found that for all eight compounds there was at least one feasible pathway to produce them with the maximum theoretical yield of 100% g-C, without loss of carbon to other byproducts.

For most of the target compounds, there exist linear pathways, identified by the graph search algorithms, that can be integrated into the host metabolism without the need of network expansion. We compared the product yield of such linear pathways with the product yield when SubNetX was used, and we found that higher yields could be achieved in the latter case. We analyzed the networks of the higher yield, and we found that they are operating as branched pathways of heterologous reactions that converge to product synthesis. These observations are illustrated via three pathway examples (Figure 3). The first pathway (Figure 3a) was linear and required four reactions to derive benzyl cinnamate from phenylalanine. As the second step of this pathway produced a byproduct not used in subsequent steps (i.e., glyoxylate), this carbon loss reduced the product yield. When we used the extracted network from SubNetX a branched pathway to benzyl cinnamate (Figure 3b) derived from two native precursors, phenylalanine and 4-hydroxybenzoate. No byproducts were formed in this pathway, and all carbon atoms were conserved to produce the target. This demonstrates that SubNetX can provide feasible pathways, which can increase the product yield by reducing byproducts, which could even be toxic due to accumulation in the host.

**Figure 3:**
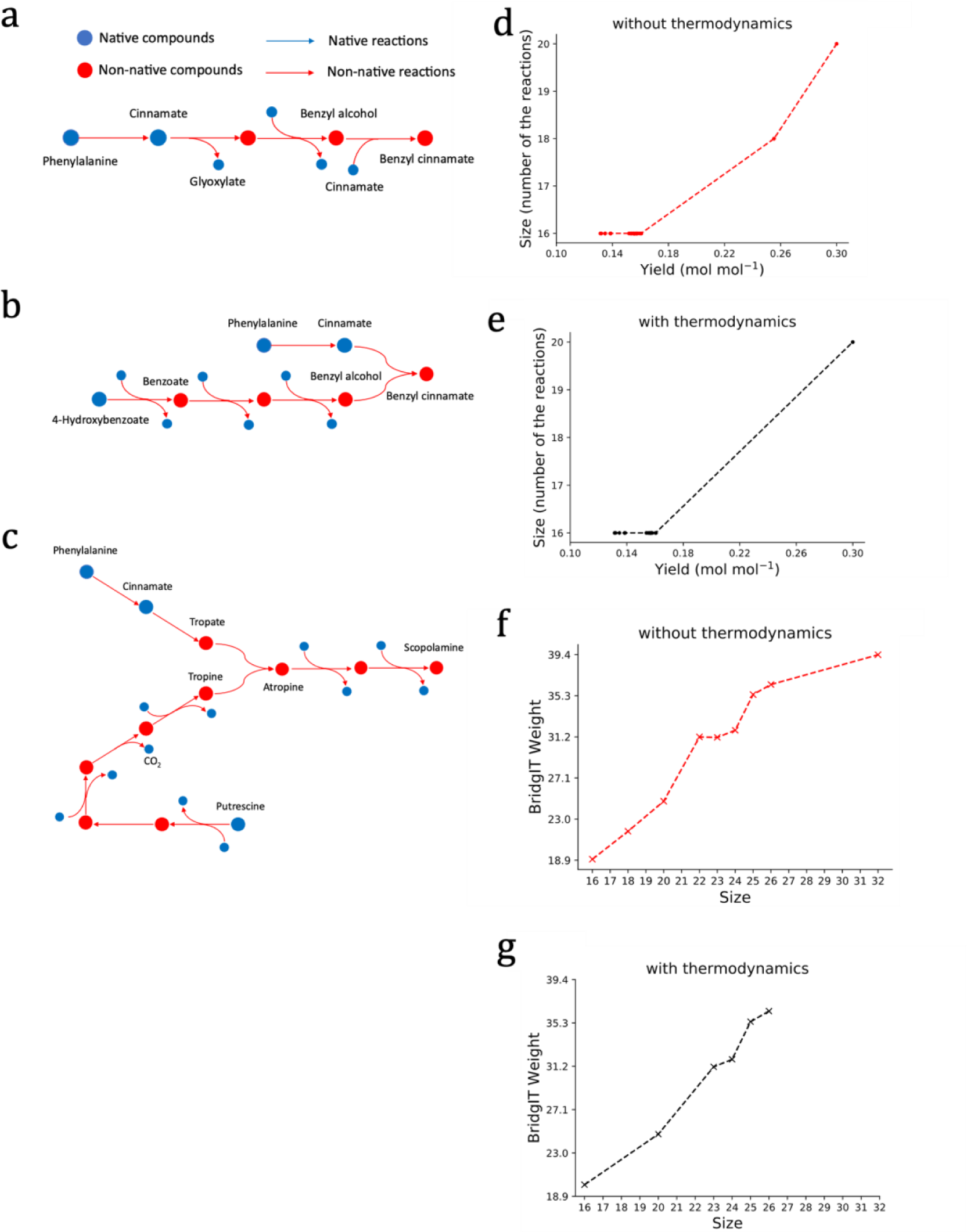
Examples of feasible pathways and exploring the trade-off between size, yield and BridgIT weight for berberine. **a** A linear pathway comprising four reactions produced benzyl cinnamate. However, the formation of glyoxylate as a byproduct, which could not be used in the subsequent steps, caused a loss of some carbon atoms. **b** While some carbon atoms were lost in linear pathways, branched pathways produced benzyl cinnamate while preserving all carbon atoms and thus had higher yields than linear pathways. **c** Scopolamine could not be produced using linear pathways, as different parts of its complex structure must be derived from different precursors. For example, in this pathway, tropine was derived from putrescine, and tropate was derived from phenylalanine. Then, tropate and tropine were combined to form atropine, which in turn formed scopolamine in two steps. **d** The Pareto fronts for berberine showed that achieving higher yields requires integrating additional reactions. **e** Integrating thermodynamics impacted the Pareto front only for berberine and no other compound (Supplementary Fig. 3). **f** Although the general trend showed that the minimum BridgIT weight increased with size, such an increase was not monotonic; in some cases, lower BridgIT weights were obtained by increasing the size. **g** Integration of thermodynamics indicated the infeasibility of some longer pathways for berberine (for other compounds, see Supplementary Fig. 4).

The third example concerned the production of scopolamine. The shortest feasible pathway toward this compound (Figure 3c) contained two separate branches of heterologous reactions to form tropate and tropine, each of which contained a part of scopolamine. Because different parts of its structure would have to be derived from different native precursors, there was no linear pathway of heterologous reactions to produce scopolamine. This example shows that SubNetX can extensively search far beyond the linear pathways obtained by the commonly used graph search algorithms.

### Predicted pathways promise higher yields than known experimentally implemented pathways

To compare the pathways predicted by SubNetX with those that have been implemented experimentally (Table 1), we first need to make the data comparable; experimentally implemented pathways are typically reported as a gene set (Supplementary Table 4), whereas the output of SubNetX is a reaction set. We, therefore, mapped the genes reported for the experimental pathways to their corresponding enzyme classes (EC) using UniProt (Supplementary Table 4) and mapped the predicted reactions of the pathways to their EC classes using BridgIT^18^.

**Table 1.** Comparison of the pathways predicted with SubNetX with the pathways retrieved from the literature.

| compound | pathway length |  | predicted<br>molar yield | # pathways<br>with better<br>yield than the<br>best-matched<br>pathway |
| --- | --- | --- | --- | --- |
|  | literature | predicted |  |  |
| Benzyl benzoate | 11 | 4–10 | 0.25–0.43 | 81 |
| Benzyl cinnamate | 6 | 4–14 | 0.22–0.38 | 45 |
| N-cinnamoyl serotonin | 5 | 5–17 | 0.24–0.32 | 71 |
| Berberine | 13 | 16–26 | 0.13–0.30 | 31 |
| Ajmalicine | 17 | 18–20 | 0.16–0.29 | 66 |
| Quercetin 3-O-(6'-acetyl-glucoside) | 9 | 7–12 | 0.21–0.26 | 0 |
| Scopolamine | 20 | 10–28 | 0.19–0.35 | 0 |
| Strictosidine | 15 | 14–22 | 0.14–0.22 | 0 |

Then, for comparison, we looked at the first three or four levels of the EC numbers to match either the enzyme family or the exact enzyme. The pathways predicted by SubNetX retained some parts of the experimentally implemented pathways. For example, the predicted pathway with the highest overlap for ajmalicin had 6 out of 12 reactions in common with the experimentally implemented pathway, matching the first three levels of the EC numbers. Interestingly, 66 pathways predicted higher yields than those with the highest overlap (Table 1). This could have several reasons, including that the predicted pathways have been optimized for *E. coli*, whereas some experimental pathways were implemented in other organisms. Another reason could be that the experimentally implemented pathways might include reactions with unknown mechanisms or stoichiometries, which were not included in the model because the SubNetX search space is limited to a database of mass-balanced reactions with known stoichiometries. Once such reaction stoichiometries and mechanisms are known, they can be easily integrated into the SubNetX search space, e.g. ARBRE. Finally, this result demonstrates the ability of SubNetX to reconstruct pathways with higher yields, although this will need to be validated experimentally. Thus, despite limitations such as incomplete stoichiometric or mechanistic information and differences in annotation, we have demonstrated that SubNetX can retrieve a subset of experimentally implemented pathways while suggesting potentially more efficient alternatives for the remaining pathways.

### Ranking the predicted pathways using constraint-based optimization

Next, we sorted the pathways by assigning them a weight either by length or enzyme specificity using the MILP formulation. To weight pathways by length, we assigned equal weights to all reactions. To weight the pathways based on the likelihood of a correct enzyme assignment, we used the BridgIT score^18^ (see Methods) to weight the reactions. Here, if the pathway contains novel reactions, a lower weight reflects a higher likelihood that promiscuous enzymes exist to catalyze these novel steps^18^.

With this weighing system, we first explored the trade-off between the yield and the pathway size by varying the lower bound for the target production to 25%, 50%, 75%, and 100% of the maximum theoretical yield, and for each of these bounds we computed the minimum pathway size. For all compounds, increasing the product yield beyond a certain threshold increased the pathway size (Figure 3d and Supplementary Fig. 3). We also investigated how thermodynamic constraints affected the trade-off between yield and size by exploring the Pareto front (Figure 3e and Supplementary Fig. 3). Here, accounting for thermodynamics did not affect the Pareto optimal front for any compound besides berberine, which had a reduced number of pathways on the Pareto optimal front due to thermodynamic infeasibilities (Figure 3e). This indicates the inclusion of thermodynamics may alter the trade-off between size and yield, and the shape of the Pareto front. In general, the inclusion of thermodynamics could reduce the optimal yield, suggesting that thermodynamics-based flux analysis^19,20^ should always be applied for more reliable predictions.

To further analyze the pathways based on BridgIT weight, we plotted the minimum BridgIT weight of each pathway against pathway size (Fig. 3f and Supplementary Fig. 4). Here, lower BridgIT weights were preferred because they represent a shorter pathway with a higher probability of an appropriate enzyme assignment. The general trend in the rankings showed that the BridgIT weight increased with size, although not monotonically, suggesting that there are some cases where better enzyme assignment can be achieved by increasing pathway size. As before, thermodynamics did not affect the trade-off between size and BridgIT weight, except for berberine (Figure 3g and Supplementary Figure 4), as seen in the rankings performed entirely by weight. Also demonstrating the flexibility of this method, the BridgIT weighting system can be modified to further refine the results according to the user’s needs by adjusting the coefficients in the objective function. For example, we adjusted the BridgIT weights to give more weight to the relevance of enzyme predictions than to the size of pathways, thus changing the correlation between BridgIT weight and size (Supplementary Figure 5).

Finally, we evaluated the fraction of thermodynamically feasible pathways for each compound. The highest fraction was found for N-cinnamoyl serotonin, where all pathways were feasible. The fraction of thermodynamically feasible pathways was greater than 90% for ajmalicine, benzyl cinnamate, scopolamine, benzyl benzoate, and quercetin 3-O-(6’-acetyl-glucoside) (92%, 95%, 97%, 95%, and 99%, respectively). For strictosidine, 90% of the pathways found were feasible, with all infeasibilities occurring on the shortest pathways (Supplementary Fig. 6). However, berberine showed a much lower feasibility, with only 55% of the pathways considered thermodynamically feasible. This is consistent with the observation of a significant change in the Pareto fronts after the integration of thermodynamics.

We further investigated the reason for such a high level of thermodynamic infeasibilities for berberine. We found that all infeasible pathways involved an identical step in which (S)-coclaurine underwent formaldehyde methylation accompanied by hydrogen peroxide reduction to water. The estimated standard free Gibbs energy for this reaction is highly positive (22.75 +/-0.03 [kcal mol^-1^]). This suggests that by removing the reactions with highly unfavorable standard free Gibbs energies, we can reduce the size of the subnetwork and avoid thermodynamically infeasible pathways. Such considerations are implemented in SubNetX and can be controlled by user-defined parameters.

To assess whether thermodynamic feasibility correlated with the other ranking metrics, we also stratified the pathways based on size or yield and calculated the fraction of feasible pathways in each stratum (Supplementary Fig. 6). The results indicated that for some compounds, the fraction of feasible pathways increased with size (e.g., ajmalicine and berberine), and for some, it increased with yield (e.g., benzyl cinnamate and berberine). However, we could not find a general correlation between thermodynamic feasibility and size or yield that applied to all cases.

### SubNetX predicts biosynthetic pathways toward complex non-natural compounds

While the price of chemically synthesized drugs can be prohibitively expensive for patients, biochemical production from simple precursors can reduce costs^5,21^. For example, tadalafil is an erectile dysfunction drug produced by chemical synthesis with no known biosynthesis pathways or enzymes in natural organisms^22^. In order for SubNetX to suggest biological alternatives for the synthesis of tadalafil, we first had to assemble the networks, ensuring that all possible reactions for its synthesis were included. Therefore, we first used SubNetX to expand a known pathway previously used for noscapine pathway derivatives^23^. This allows us to expand the biochemical search space so that we do not miss potential alternative pathways with higher yields than the known pathway. We also selected every intermediate in the organic synthesis pathway of tadalafil^24^ and extended the network of predicted biochemical reactions to include ones for their synthesis using BNICE.ch, which reconstructs known biochemical reactions as well as generates novel, hypothetical reactions^25^. We added this network to ARBRE to expand the SubNetX search space.

We next used SubNetX to find possible subnetworks for synthesizing tadalafil. To simultaneously determine the impact of adding flexibility to the algorithm in terms of the number of reactions for the synthesis of cosubstrates, we applied SubNetX with three different settings, selecting only the pathways with the a) minimal number of reaction steps required to connect cosubstrates to the host, b) minimal+1 steps, and c) minimal+2 steps. To analyze the potential for successful integration of the extracted subnetworks with the host model, we applied topological network analysis (Table 2), and we evaluated whether tadalafil was included in the main component of the network, indicating its connection to the host metabolism. The minimal and minimal+1-step pathways limited the subnetwork to only 36 reactions and kept tadalafil out of the main component of the subnetwork. Increasing the size of pathways to minimal+2 extracted a subnetwork of 10,611 reactions with tadalafil integrated into the main component, indicating a potentially successful integration with the host model.

**Table 2.** Subnetwork extraction for tadalafil, showing the number of compounds and reactions in the extracted subnetworks for the three settings.

| Metric | Minimal PW | Minimal + 1 | Minimal + 2 |
| --- | --- | --- | --- |
| # compounds | 17 | 24 | 2,986 |
| # reactions | 21 | 36 | 10,611 |
| # network edges | 16 | 26 | 6,198 |
| # isolated network components | 3 | 3 | 1 |
| % compounds in the main component | 53% | 67% | 100% |
| Is the target compound in the main component? | No | No | Yes |

We then evaluated the three extracted subnetworks for yield and feasibility. For the minimal and minimal+1 steps, the model predicted no tadalafil production because the connection between the cosubstrates and the host metabolism was not properly established. In contrast, the minimal+2 steps subnetwork produced tadalafil with the maximum theoretical yield. We then used this subnetwork to investigate the trade-off between yield and size, as described above for the example syntheses. As with the natural compounds, including more reactions allowed for higher product yields (Figure 4c). Finding pathways for the synthesis of tadalafil in *E. coli* demonstrates that SubNetX can suggest biosynthesis pathways to produce non-natural compounds for which natural production pathways are not known.

**Figure 4:**
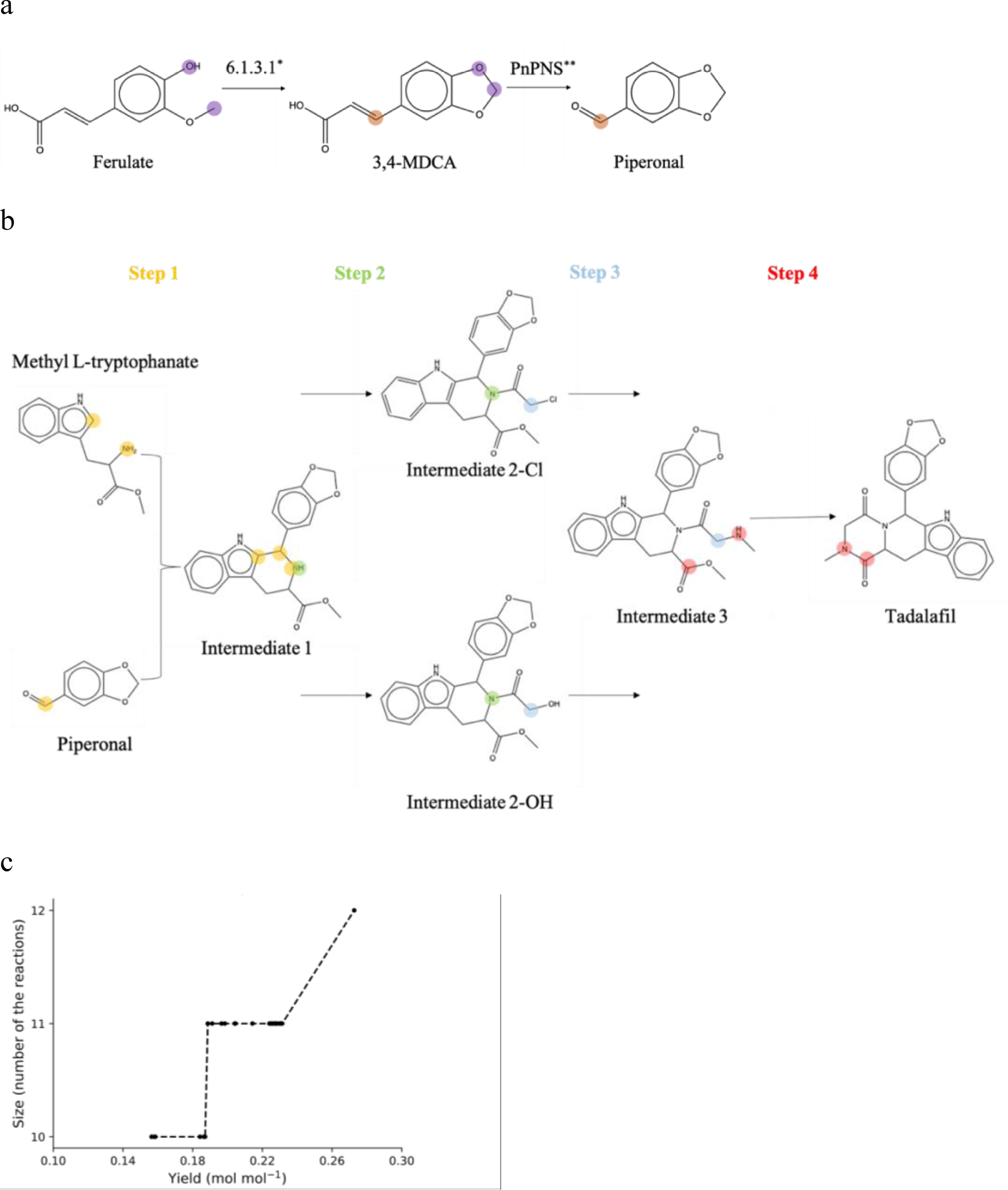
Pathway prediction for the biosynthesis of tadalafil. **a** Tadalafil synthesis pathway used for BNICE.ch network expansion. Atoms of the reactive site are highlighted. Biosynthesis route of piperonal, a precursor of tadalafil (adapted from *Jin et al.*^39^, *enzyme proposed in this work, **enzyme proposed by *Jin et al.*). 3,4-MDCA: 3,4-methylenedioxycinnamic acid. **b** Chemical synthesis of tadalafil, adapted from Baumann et al.^24^ (Step 1) and expanded with necessary intermediates (Steps 2 and 3). The upper path corresponds to the chemical synthesis of Intermediate 2-Cl, and the lower to the proposed biological alternative, Intermediate 2-OH. **c** Exploring the trade-off between size and yield for tadalafil.

## Conclusion

We present SubNetX, a method for exploring large networks of reactions to find an optimal pathway for the synthesis of a target compound. An important feature of our method is the integration of two previously independent approaches to pathway design: graph-based pathway search and constraint-based optimization. The linear pathway search algorithms easily explore large biochemical networks (e.g., ARBRE and ATLASx), and the constraint-based optimization finds the most relevant pathways in terms of yield and feasibility. A second important and novel feature of SubNetX is the extraction of a balanced subnetwork of reactions around the linear pathways from the graph search, which can be seamlessly integrated into the genome-scale metabolic model of any organism. Using the SubNetX workflow, we were able to predict pathways for the synthesis of natural products of varying complexity. This work showed that the assembly of multiple reactions into networks from multiple native precursors that converge to product synthesis is necessary for several classes of secondary metabolites, and such pathways could improve production yields and reduce potentially toxic by-products. We also demonstrated that SubNetX can retrieve experimentally implemented pathways and suggest potentially more efficient alternatives, even in the absence of stoichiometric information or annotation differences. Furthermore, application of SubNetX to a complex non-natural therapeutic, tadalafil, demonstrated its ability to predict biosynthetic pathways for compounds not produced by any known organism.

Experimental investigation and validation of the pathways proposed by SubNetX will allow for its immediate use in a variety of organisms and any application requiring the design and analysis of metabolic pathways. For instance, though our included examples were all in *E. coli*, SubNetX can be readily extended to other host organisms containing a genome-scale model with metabolites annotated using appropriate identifiers. Furthermore, SubNetX is not limited to pathway design for the production of complex molecules and can be used for other applications, such as the biodegradation of xenobiotics^26^, designing a synthetic consortium of microorganisms with emergent properties^27,28^, or analyzing the interactions between hosts and parasites^29^. Finally, SubNetX is compatible with more advanced constraint-based methods for strain engineering^30–33^, allowing the optimization of chassis organisms for industrial applications. We envision that SubNetX will aid in a variety of strain-design optimizations due to its ability to evaluate many possibilities and rank the best solutions based on user-defined criteria.

## Methods

### Selection of the case study compounds

We selected the case studies based on two criteria. First, they should be present in the ARBRE network. Second, their experimental biosynthesis should be available in the literature to compare the results (Supplementary Table 3, Supplementary Table 4). The synthetic accessibility (SA) score was calculated using the rdkit implementation of the SA score^14^.

### Reaction network data

Recently, we generated the largest known network of predicted biochemical reactions, ATLASx^11^. We showed that it is possible to translate reaction networks into graphs connecting at most two compounds per edge and use computational algorithms to navigate these graphs efficiently^34^. In another work, we presented ARBRE, demonstrating that additional data curation and user-defined parameters might refine the pathway ranking to propose only the most relevant pathways as a starting point for experimental tests^13^. ATLASx and ARBRE integrated the reactions into the reactant-product pair (RP-pair) network. This work used ARBRE as the reaction database to search for the pathways. In the future, the search space for the reactions can be expanded to ATLASx reactions or any other set of balanced reactions.

### Preparing the search space for pathways

We obtained a set of elementally balanced reactions, both known and novel, from ARBRE^13^. For each RP-pair in the network, we attributed all the associated reactions and identified the sets of boundary metabolites. The initial pathway search and the network expansion used the RP-pairs, corresponding reactions, and boundary metabolites. The reactions with their corresponding stoichiometry were stored and integrated into the GEM.

### Translating reaction network into an explorable hypergraph-like network

The best representation of the structure of biochemical networks is through hypergraphs. However, algorithms to navigate hypergraphs are not as developed as those to predict linear pathways within a simple graph^2,35^. Simple graphs and linear pathway search algorithms allow assigning weights to graph edges based on atom conservation so that pathways with higher theoretical yields are prioritized^34^. However, while searching for linear pathways, there is no guarantee that the cosubstrates of reactions are also connected to the host metabolism. To preserve the information about all the substrates and products of a reaction within a single edge, as in hypergraphs, we developed a method to integrate cosubstrate information into a simple graph as an edge property. In addition, we introduced a filtering step to exclude unbalanced reactions to avoid problems in pathway evaluation, as unbalanced reactions can produce chemicals without consuming any resources.

### Integrating host metabolites with the reaction network

Host metabolites are regularly annotated in models by public identifiers. We mapped the host metabolites to their corresponding atom connectivity structure without considering cellular compartments. Metabolites without a defined molecular structure were excluded. This allowed us to map the host metabolites to the compounds in ARBRE. We collected the list of metabolites in *E. coli* from iJO1366^17^. 800 of 1099 native metabolites in *E. coli* were mapped to the compounds in ARBRE using the molecular structure. The mapped metabolites then served as precursors for linear pathways.

### Selecting host organism

SubNetX is applicable to all organisms with available GEMs. To apply SubNetX to other organisms, it is sufficient to map the new host metabolites to the identifiers of the search space of choice, e.g., ARBRE. Here, we applied this pipeline to *E. coli* using the metabolic model iJO1366.

### Controlling the properties of the extracted subnetwork

We have defined a set of parameters that allow users to control the size and coverage of the subnetwork. The user can define the precursor or select from the host native metabolites. Precursor filtering parameters allow for the specification of whether a substructure filtering should be applied. These parameters also define the number of similar precursors to select. A network filtering parameter specifies the threshold for conserved atom ratio per each pathway step. Users can define the number of linear pathways in the core set and the number of linear pathways to connect cosubstrates. Moreover, there is the option to select only the shortest or the shortest+X-step pathways to connect the cosubstrates, where X is the number of additional steps to include. Users can indicate their preference to include known reactions and longer pathways using the exponential transformation of the graph distances^34^.

### Filtering the precursor set based on compound structure

The user-defined set of precursors can be filtered based on chemical substructures. To this end, the user can introduce a *.mol* file describing the desired substructure that should be present in all precursors. We used the rdkit.Chem library (https://www.rdkit.org) to match the substructures. The output of this step is a list of precursors corresponding to the desired structure. The precursors with more carbon atoms than the target (either the primary target or cosubstrates) are removed to achieve an optimal carbon flow. For the remaining precursors, the rdkit.Chem.rdFMCS library is used to calculate the maximum common substructure. The *N* compounds (a user-defined parameter) with the maximum number of atoms in the common substructure are then selected.

### Filtering the network based on user-defined parameters

The network is prepared for the pathway search as described elsewhere^13^. The unwanted compounds and RP-pairs can be removed from the network based on molecular structures. This network filtering ensures that the pathway search and network expansion avoid unwanted compounds and use only the reactions with desired properties.

### Finding the core pathway set

The initial linear pathways, subsequently called the core pathway set, consist of linear pathways connecting the target compound to the defined precursors (as described in NICEpath^34^ and ARBRE^13^). Such pathways connect the precursors defined in the first step to the target compound. The pathways are found using the k-shortest loopless path algorithm implemented in the networkx library (https://networkx.org). The pathway search algorithm takes user-defined parameters to determine the total number of pathways and the maximum pathway length.

### Subnetwork expansion

Subnetwork expansion is the central step of the pipeline, connecting all the atoms of the target compound to the host metabolism. The initial subnetwork consists of the core pathway set. We associate cosubstrates with the edges of the subnetwork, where each edge represents a step of the pathway. We find linear pathways to connect the non-native metabolites to the host to ensure that all cosubstrates can be produced and all byproducts can be consumed. The new pathways are added to the subnetwork, and new boundary metabolites are specified. The expansion continues until non-native metabolites are connected to the host metabolism. In particular cases, non-native metabolites cannot be connected to the host due to the absence of the pathway in the initial network (i.e., ARBRE) and must be removed in the subnetwork convergence step. We search for reactions with the fewest cosubstrates outside the host metabolism to avoid expanding the subnetwork toward the non-natural cofactor pairs. For each cosubstrate, precursors are selected as described above. If no pathway to the host can be found, the cosubstrate is omitted, and the pathways relying on it will be excluded during the subnetwork convergence. The expansion terminates once all the cosubstrates are either connected to the native metabolites or omitted.

### Subnetwork convergence

We remove the non-native metabolites not connected to the host metabolism during the subnetwork convergence. For this purpose, we remove all the cosubstrates not integrated into the subnetwork and the reactions that require these cosubstrates. Removing these reactions from the subnetwork might disconnect other cosubstrates from the host metabolism. This step is repeated until no cosubstrate out of the subnetwork is left. Thus, we generate a network with a bioproduction or biodegradation pathway for any boundary metabolite. We did not connect metal cofactors, e.g., [4Fe-4S] iron-sulfur cluster, Na^+^, Co^2+^, Mg^2+^, Cu^2+^, and certain gases, e.g., hydrogen selenide, nitrogen, H2, to the host if they were cosubstrates. The reactions and compounds of the converged network are the output that integrates into the host metabolism for further evaluation and ranking.

### Graph visualization

The extracted subnetwork is visualized automatically using the networkx library. Graphs in Gephi format output (.gdf) are generated, which can be imported into Gephi for visualization and further analysis.

### Integration of the subnetwork into the host model

We need to integrate the subnetwork into the host’s GEM to find feasible pathways. For this purpose, the metabolites and reactions in the subnetwork should be mapped to their counterparts in the GEM. Then, we construct an integrated network that includes all native and non-native metabolites and reactions. A demand reaction is added for the target to take this compound out of the cytosol, and we set a lower bound for this reaction to ensure that the target is produced. We used the identifiers from our internal database to map the metabolites and reactions. However, other identifiers can be used alternatively if the metabolites and reactions of the GEM are annotated with the same identifiers. Note that the integration quality depends on the coverage of the annotations, and identifiers with higher coverage are preferred. The integrated subnetwork can be saved in JavaScript Object Notation (JSON) format. JSON is a standard hierarchical format to keep structured data. Since many software packages, such as COBRA^36^, support this format, the saved files can be integrated into other pipelines and software for further analyses.

### Pathway search and evaluation

The next step in SubNetX is to search for feasible pathways. First, a Flux Balance Analysis (FBA) problem is solved, where the rows and columns of the stoichiometric matrix correspond to the integrated network’s metabolites and reactions, and the objective function is to maximize the target production. If the maximum flux of the production is nonzero, at least one feasible pathway exists in the network to produce the target compound. This way, we can also find the maximum production rate of the target 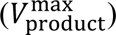 for a specified substrate uptake rate. Then, we search for and enumerate the feasible pathways by formulating a MILP problem:

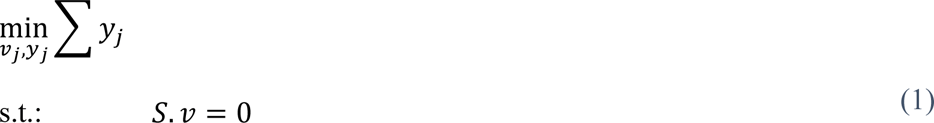

[math2

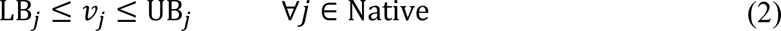

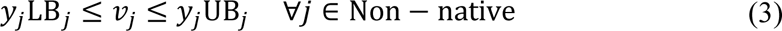

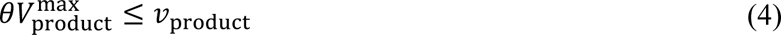

Here, a binary variable *y*_*j*_ is associated with each non-native reaction. If *y*_*i*_ is zero, the reaction is inactive and not included in the pathway. The objective function is to minimize the number of active non-native reactions. This way, we prioritize feasible pathways with the minimum number of interventions in the host. To ensure that the target compound is produced, Equation (4) sets a lower bound for the production. This lower bound is set as a fraction (θ) of 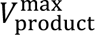 found in the previous step. We can explore the trade-off between yield and size of the pathway by adjusting θ. While higher values of θ enforce finding pathways with higher yields, lower values of θ relax the lower bound and allow us to find shorter pathways.

Integer cut constraints were added to find alternative solutions, including other optimal and suboptimal pathways:

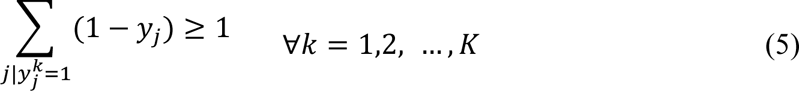

where *K* is the total number of pathways found previously and 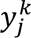 represent if the *k*th pathway includes reaction *j* or not.

### Ranking feasible pathways

In addition to the saved model as a JSON file, we save a set of tables that contain information about the feasible pathways found by the MILP formulation. These feasible pathways are ranked based on three main criteria. The first criterion is product yield, which indicates how many moles of the target can be produced per consuming one mole of the carbon source, e.g., glucose. Higher yields imply that more carbon is conserved to produce the target.

The second criterion is the weight of the pathway. The weight collectively refers to any type of preference that can be quantified and associated with the reactions. The weight of a pathway is the sum of the scores of its constituting reactions, where pathways with lower weights are preferred. In the simplest case, where all the reactions are weighted equally, minimizing the weight is equivalent to finding pathways with the minimum size. However, we can bias the pathways toward including or not including specific reactions by assigning different weights (e.g., based on toxicity or enzyme specificity).

Finally, the third criterion is thermodynamic feasibility. Some pathways cannot produce the target after integrating thermodynamic constraints due to the changes in reaction directionalities. The output of the thermodynamic evaluation is a binary assignment; the pathways are either thermodynamically feasible or infeasible.

### Pathway weight calculation

Various properties, such as enzyme availability or toxicity of the byproducts, can be quantified as weights and associated with the reactions. Similarly, the weight of a pathway is the sum of the weights of its constituting reactions. In the simplest case, when an equal weight is assigned to all the reactions, minimizing the weight of the pathway is equivalent to minimizing the size. However, we can prioritize including or excluding specific reactions by assigning different weights. For example, we assigned weights to the reactions based on enzyme availability. In particular, we used BridgIT scores, where higher BridgIT scores indicate a higher likelihood of finding an appropriate enzyme. We defined the BridgIT weights as follows:

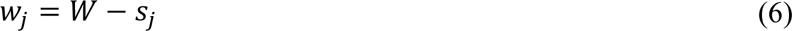

where *K*_*i*_ is the BridgIT score for the *j*th reaction. *W* is a user-defined parameter, where higher values of *W* prioritize smaller pathways, and lower values of *W* prioritize integrating reactions with higher BridgIT scores. We minimized the sum of weights of the active reactions, i.e., 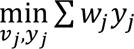. A reaction’s weight contributes to the objective function only if *y*_*j*_ = 1, i.e., the reaction is part of the pathway. We set *W* to two different values in our analyses, 1.1 and 2, to evaluate the impact of changing *W*.

### Thermodynamic evaluation

We also check the thermodynamic feasibility of the found pathways. To this end, we integrate each pathway into the GEM individually and solve Thermodynamic Flux Analysis (TFA)^19^. Like FBA, TFA optimizes an objective function subject to mass balances. In addition, TFA applies additional constraints so that the directionality of each reaction is determined based on its Gibbs free energy. The Gibbs free energy of a reaction depends on its standard Gibbs free energy, the thermodynamic properties of the environment, and the metabolite concentrations. The latter is approximated based on the available ranges for the metabolite concentrations if metabolomics data is not available^19^. The standard Gibbs free energy of a reaction can be calculated as the difference between the sum of the standard Gibbs free energy of the formation of the products and reactants.

The standard Gibbs free energies of formation for the native compounds were already calculated^37^. We used the Group Contribution Method (GCM) to estimate the standard Gibbs free energy of formation for non-native compounds^37^. We first obtained the Simplified Molecular Input Line Entry System (SMILES) for these compounds. Then, we transformed the SMILES into their major protonation state at standard pH and generated the MDL Molfiles using Marvin (version 18.1, 2018, ChemAxon http://www.chemaxon.com). The Molfiles were used in GCM to estimate the standard Gibbs free energies of formation. Since the cellular thermodynamic properties such as pH and ionic strength are different from the standard condition, we corrected the standard Gibbs free energies of the formation using the Debye-Hükel approximation. Finally, the corrected Gibbs free energies of formation and the metabolite concentrations were used to calculate the Gibbs free energies of the reactions^38^.

### Preparing the database to find pathways for tadalafil

For the first and fourth steps of the proposed biosynthesis pathway of tadalafil (Figure 4a), we added two new BNICE.ch reaction rules based on EC classes 4.2.1.78 (norcoclaurine synthase) and 4.2.1.145 (capreomycidine synthase), respectively. We identified that the second and third steps could be catalyzed by the reverse reaction of EC 3.4.17.23 (angiotensin-converting enzyme 2) and by 6.3.4.12 (glutamate-methylamine ligase), respectively. One of the precursors of the chemical synthesis of tadalafil, piperonal, is a natural compound, though it does not have a fully characterized biosynthesis route that would connect it to any of the host metabolites in the ARBRE network. To address the biosynthesis of piperonal, we started with the precursor ferulic acid. From this, we added new BNICE.ch rules based on 6.1.3.1 (olefin β-lactone synthetase) to convert ferulic acid into 3,4-methylenedioxycinnamic acid (3,4-MDCA) and the gene PnPNS^39^ to convert 3,4-MDCA into piperonal (Figure 4b). In all, expanding the BNICE.ch network and identifying all the reactions carrying flux from tryptophan, as the native precursor, toward tadalafil added eight auxiliary reactions to the ARBRE network. This ensured that all the necessary reactions were present to reconstruct a balanced subnetwork to synthesize tadalafil.

## Author Contribution

AS, OO, and VH designed and conceptualized the study and developed the methodology. AS implemented the software and ran the simulations to extract subnetworks. OO implemented the software and ran the simulations to evaluate and rank the pathways. AS validated the predicted pathways with the reported pathways in the literature. AS and OO curated the data, developed the visualizations, performed formal analysis, conducted the investigation, and wrote the original draft. AS, OO, and VH discussed and analyzed the results. AS, OO, and VH wrote and reviewed the manuscript. VH was responsible for acquiring resources, funding, project administration, and supervision.

## Code and Data Availability

Subnetwork extraction and analysis code and data are available at https://github.com/EPFL-LCSB/SubNetX.

## Acknowledgment

We would like to thank *Dr. Kaycie Butler* and *Dr. Ljubisa Miskovic* for their valuable comments on the language and structure of the manuscript. This work was funded by the Swiss National Science Foundation (SNSF) under grant 200021_188623, by the European Union’s Horizon 2020 Research and Innovation Program under grant agreement no. 814408, and by the École Polytechnique Fédérale de Lausanne.

