## Supplementary Information for "Designing pathways for bioproducing complex chemicals by combining tools for pathway extraction and ranking"

**Supplementary Figures**

| a  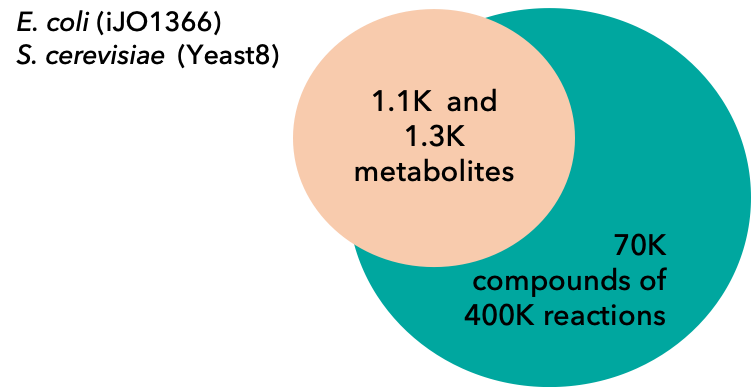  b  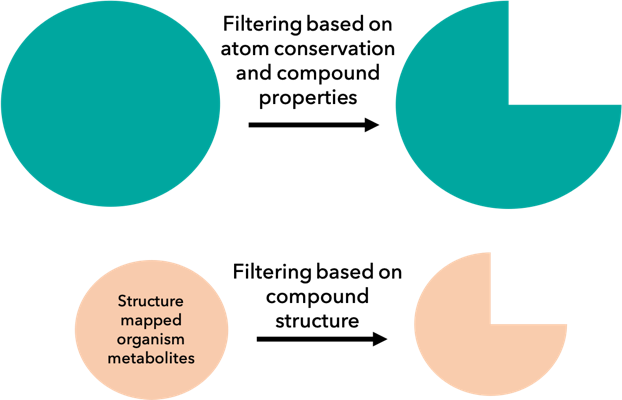  ARBRE network  c  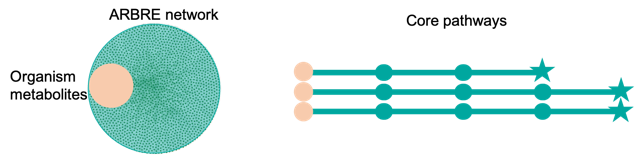 | d  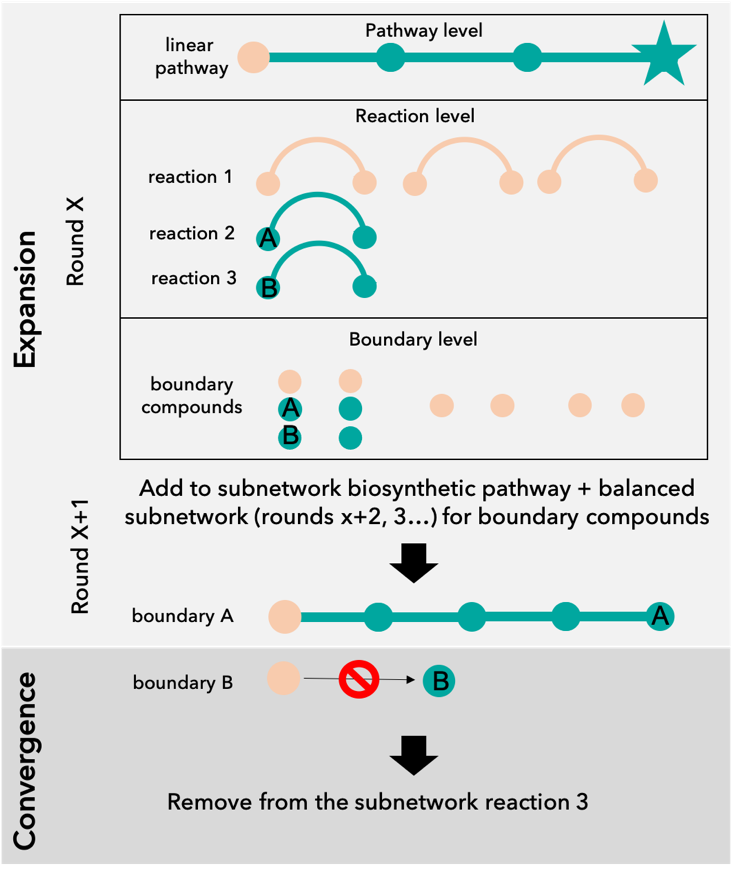 |
| --- | --- |
| 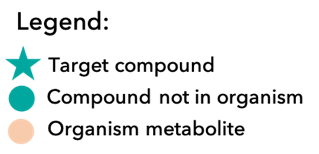 | |

Supplementary Figure 1: **Network extraction pipeline.** The subnetwork extraction is done in 5 steps (Figure 1): 1) Filtering the precursor set based on the defined compound substructure and similarity to the target; 2) Filtering the network based on user-defined parameters; 3) Finding the core pathway set; 4) Subnetwork expansion; 5) Subnetwork convergence. The subnetwork extraction algorithm allows searching for a compiled set of linear pathways connecting the target to the host metabolism through a simple graph. **a** Mapping the organism metabolites with the ARBRE reaction network; **b** Filtering based on the organism metabolite properties and network properties; **c** Predicting alternative core linear pathways towards the target metabolite; **d** Network expansion and convergence: linear pathway search and identification of pathway boundaries.

| a | c  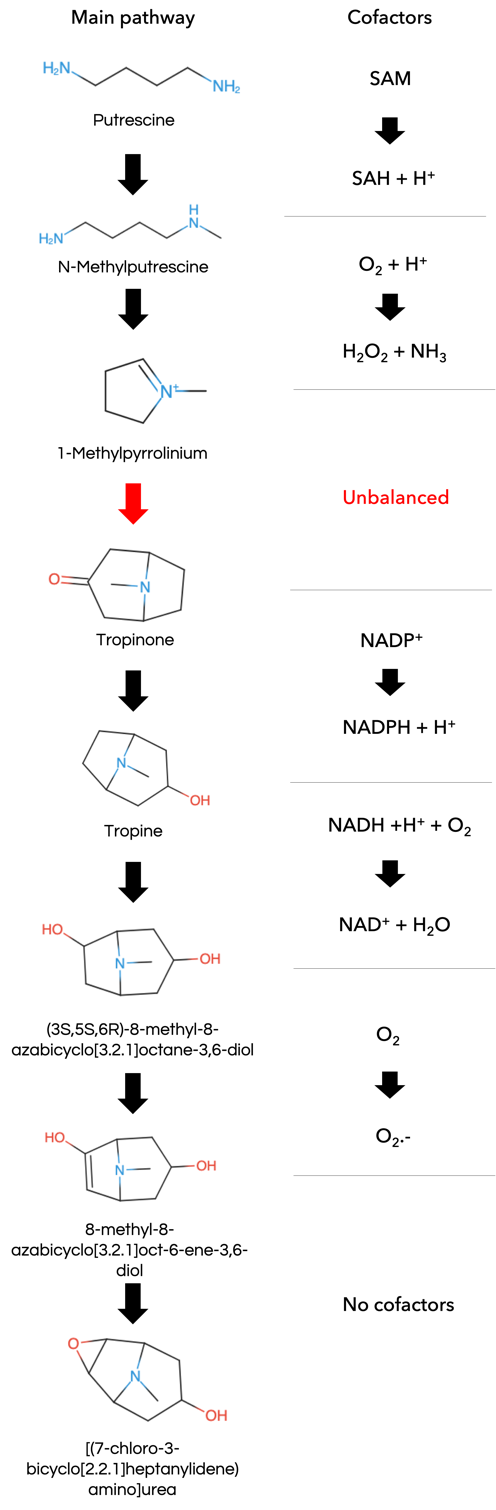 |
| --- | --- |
| LCSB ID: 59924115  (3S,5S,6R)-8-methyl-8-azabicyclo[3.2.1]octane-3,6-diol  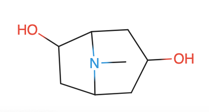  LCSB ID: 3817731  [(7-chloro-3-bicyclo[2.2.1]heptanylidene)amino]urea  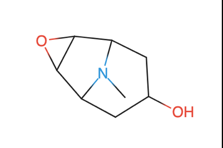 |  |
| b  Step 1  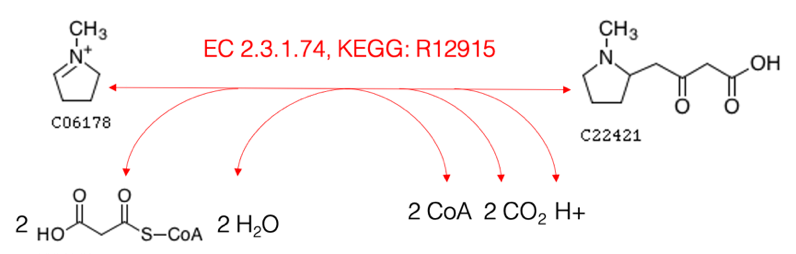  Step 2  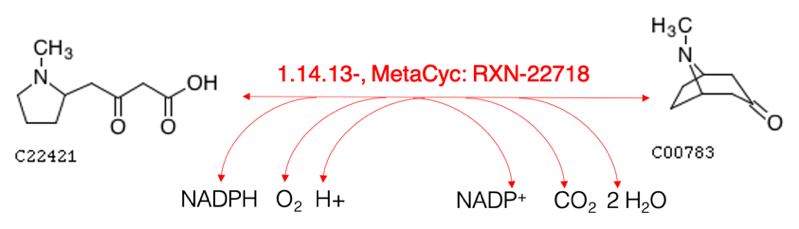 |  |

Supplementary Figure 2: **Reconstructing putrescine subnetwork** **a** 2 boundary compounds for which additional pathways from putrescine had to be added to ARBRE network to predict balanced subnetwork for scopolamine production from ATLASx network; **b** Two-step reconstruction of the unbalanced reaction in the pathway added from ATLASx; **c** Pathway added from ATLASx.

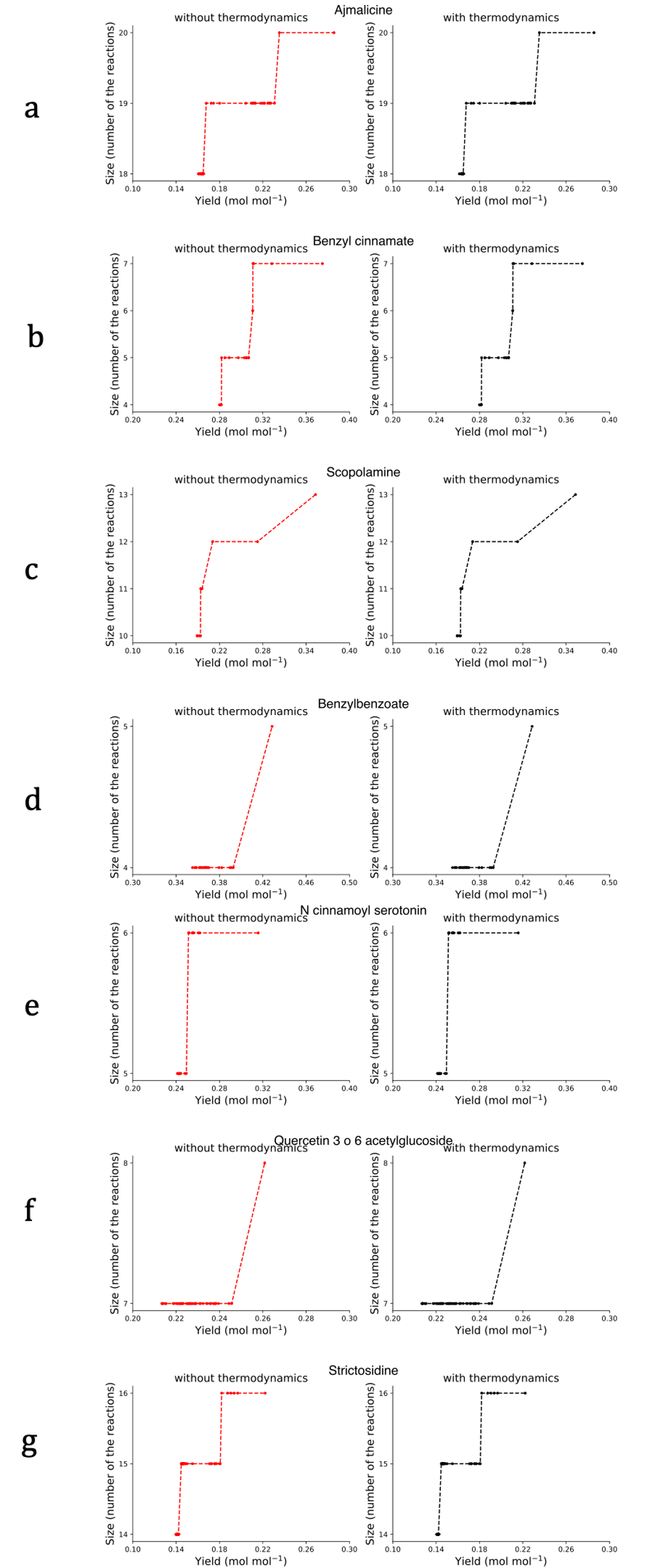

Supplementary Figure 3: **Exploring the trade-off between size and yield. a** ajmalicine **b** benzyl cinnamate, **c** scopolamine, **d** benzylbenzoate, **e** N-cynnamoyl serotonin, **f** quercetin-3-6-acetylglucoside, and **g** strictosidine. Exploring the Pareto fronts generally showed that achieving higher yields requires integrating additional reactions. Integrating thermodynamics impacted the Pareto front only for berberine, while the other compounds showed no difference.

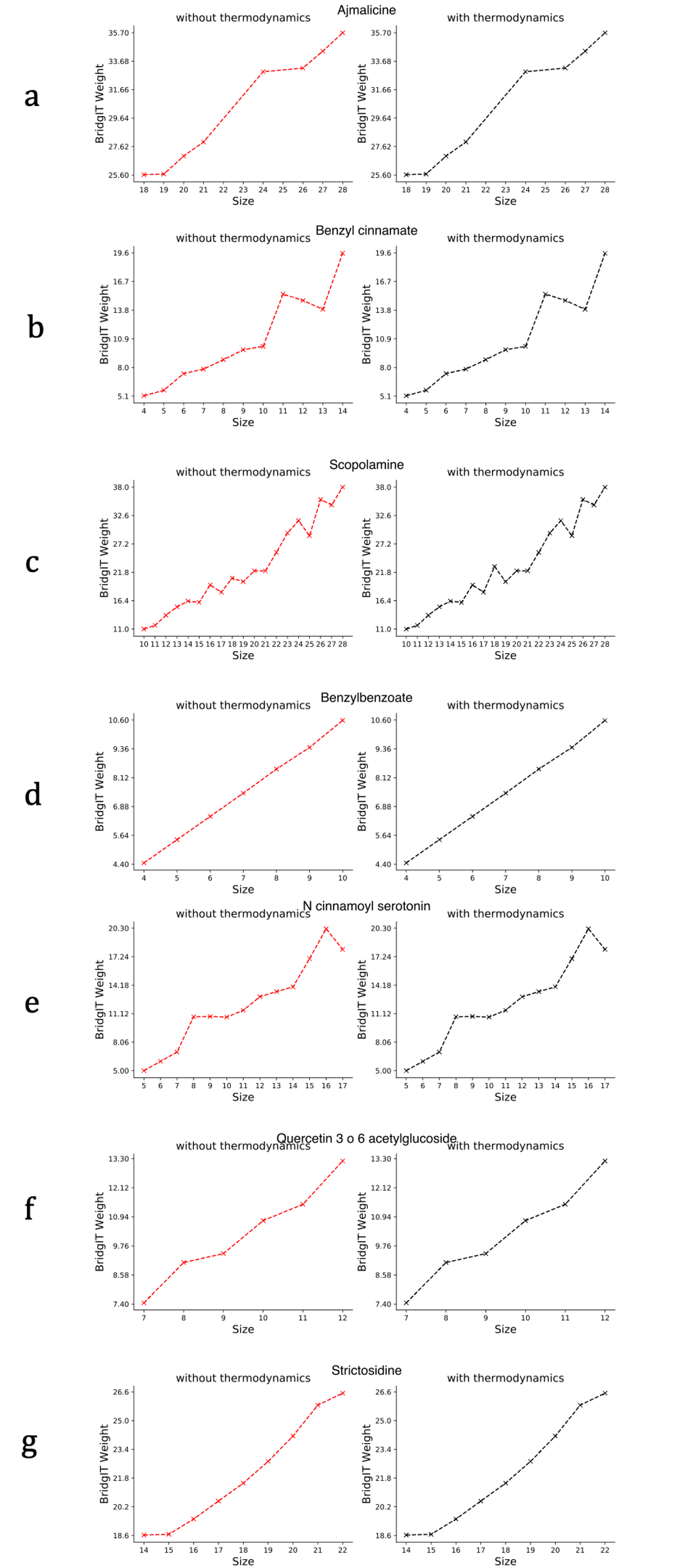

Supplementary Figure 4: **Exploring the trade-off between size and BridgIT weight. a** ajmalicine **b** benzyl cinnamate, **c** scopolamine, **d** benzylbenzoate, **e** N-cynnamoyl serotonin, **f** quercetin-3-6-acetylglucoside, and **g** strictosidine. Although the general trend showed that the minimum BridgIT weight increased with size, such an increase was not monotonic; in some cases, lower BridgIT weights were obtained by increasing the size.

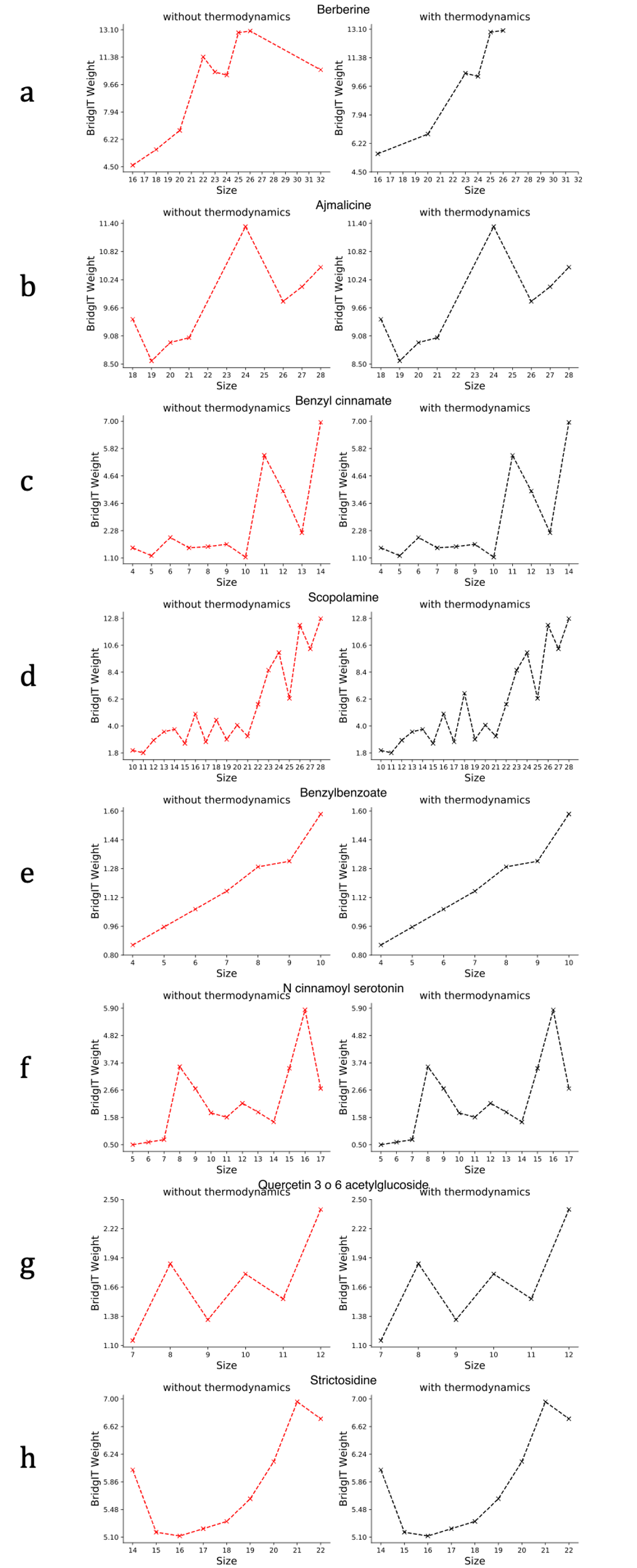

Supplementary Figure 5: **Exploring the trade-off between size and BridgIT weight with higher importance assigned to enzyme assignments. A** berberine, **b** ajmalicine, **c** benzyl cinnamate, **d** scopolamine, **e** benzylbenzoate, **f** N-cynnamoyl serotonin, **g** quercetin-3-6-acetylglucoside, and **h** strictosidine. BridgIT weight is defined as a function of size and BridgIT score, where the latter indicates the quality of enzyme assignments. Here, we assigned a higher weight to BridgIT scores to prioritize finding relevant enzymes. With the new settings, larger pathways had lower BridgIT weights due to prioritizing better enzyme assignments.

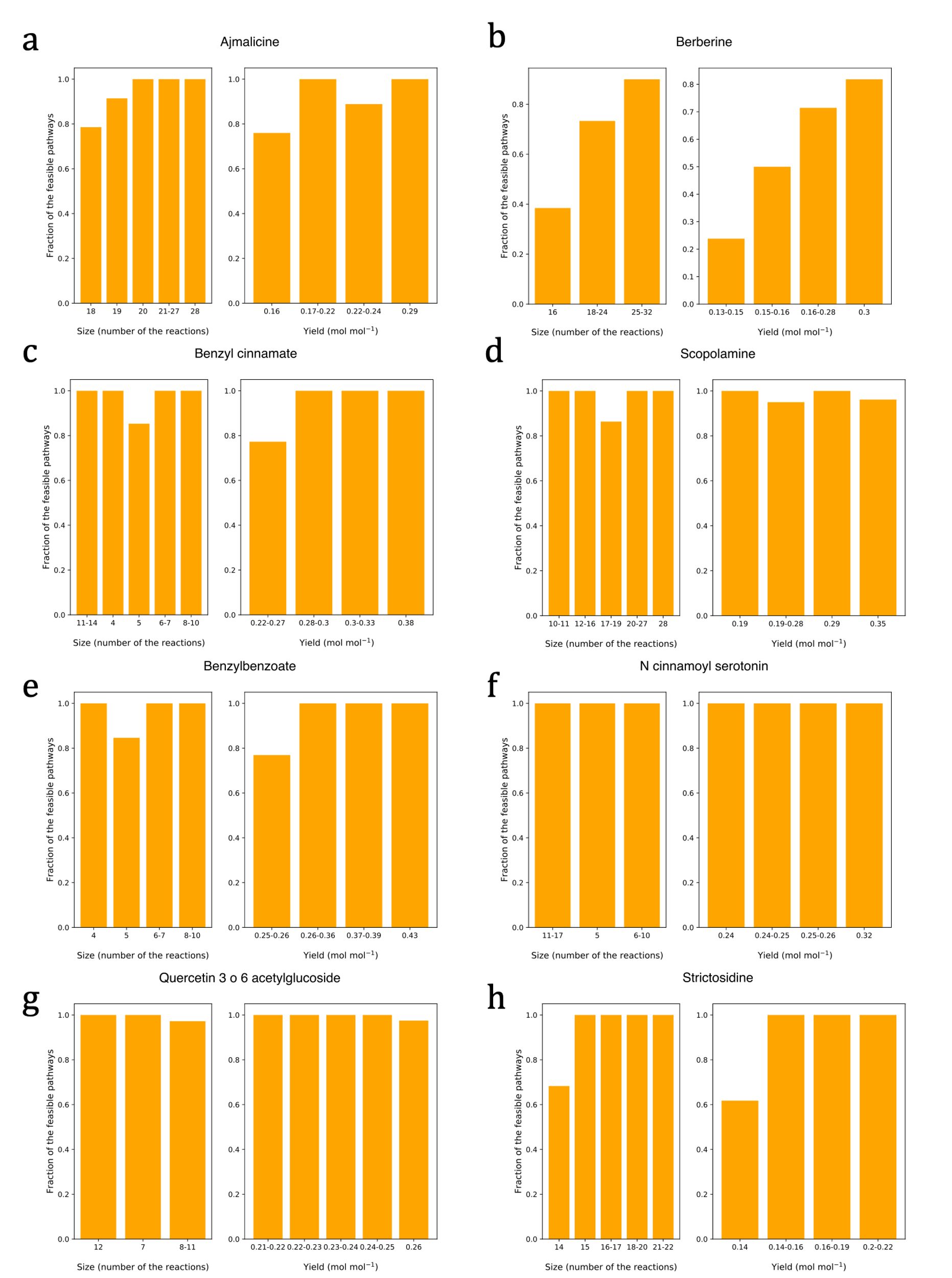

Supplementary Figure 6: **Fraction of thermodynamically feasible pathways for different sizes and yields. a** ajmalicine, **b** berberine, **c** benzyl cinnamate, **d** scopolamine, **e** benzylbenzoate, **f** N-cynnamoyl serotonin, **g** quercetin-3-6-acetylglucoside, and **h** strictosidine. We did not observe a global trend between thermodynamic feasibility and size or yield. However, compound-specific patterns were observed; thermodynamic feasibility increased with size for ajmalicine and berberine and with yield for benzyl cinnamate and berberine.

**Supplementary Tables**

Supplementary Table 1: Case study compounds tested for subnetwork extraction and pathway search. The number of compounds and reactions in the extracted subnetworks for the eight natural case study compounds.

| **Compound name** | **SA^*^ score** | **number of compounds** | **number of reactions** |
| --- | --- | --- | --- |
| Benzyl benzoate | 1.21 | 8'366 | 33'579 |
| Benzyl cinnamate | 1.54 | 8'402 | 33'989 |
| Cinnamoyl serotonin | 2.10 | 8'362 | 33'656 |
| Berberine | 2.79 | 8'464 | 34'183 |
| Ajmalicine | 3.94 | 8'390 | 33'852 |
| Quercetin 3-O-(6'-acetyl-glucoside) | 4.08 | 8'434 | 34'122 |
| Scopolamine** | 4.64 | 8'491 | 34'410 |
| Strictosidine | 4.85 | 8'389 | 33'951 |

* SA – synthetic accessibility

** After adding the required reactions from the ATLASx and the latest versions of the KEGG and MetaCyc databases

Supplementary Table 2. Small biological molecules that have already been produced in a host organism have synthetic accessibility (SA) score^14^ between 1 (easy to make) and 6.5 (difficult to make).

| **Class of compounds** | **Examples (SA score)** |
| --- | --- |
| alkaloids | Berberine (2.79), ajmalicine (3.94), scopolamine (4.64), strictosidine (4.85), staurosporine (4.95) |
| products of shikimate pathway | cinnamoyl serotonin (2.10) |
| polyphenols | rosmarinic acid (2.90), curcumin (2.43), resveratrol (2.11) |
| glycosylated isoflavonoids | quercetin 3-O-N-acetylglucosamine (4.08) |
| oligomeric products of flavonoids | Proanthocyanidins (4.77) |
| lactam antibiotics | ansalactam C (6.5), carbapenem (3.85) |
| beta-lactones | salinosporamide A (4.82) |
| stilbenes | Pterostilbene (1.89) |
| fungal indole diterpenes | aflatrem 4 (6.27) |
| phosphonates | Fosfazinomycin (4.3) |
| natural colorants | Betanin (5.36), violacein (3.04) |

Supplementary table 3. Comparison of the EC classes match for the pathways predicted with SubNetX with the pathways retrieved from the literature.

|  | # three-level ECs | | # four-level ECs | |
| --- | --- | --- | --- | --- |
| compound | recovered from literature | matched | recovered from literature | matched |
| Benzyl benzoate | 8 | 0-4 | 8 | 0-2 |
| Benzyl cinnamate | 6 | 2-6 | 6 | 0-3 |
| Cinnamoyl serotonin | 5 | 2-4 | 5 | 0-4 |
| Berberine | 10 | 5-8 | 17 | 3-8 |
| Ajmalicine | 12 | 4-6 | 14 | 1-3 |
| Quercetin 3-O-(6’-acetyl-glucoside) | 9 | 1-6 | 10 | 0-5 |
| Scopolamine | 13 | 6-9 | 18 | 4-6 |
| Strictosidine | 12 | 4-6 | 15 | 1-2 |
